# A computational modeling approach for predicting multicell patterns based on signaling-induced differential adhesion

**DOI:** 10.1101/2021.08.05.455232

**Authors:** Nikita Sivakumar, Helen V. Warner, Shayn M. Peirce, Matthew J. Lazzara

## Abstract

Differential adhesion within cell populations enables the emergence of unique patterns in heterogeneous multicellular systems. This process has been previously explored using synthetically engineered heterogenous multicell spheroid systems, in which cell subpopulations engage in bidirectional intercellular signaling to regulate the expression of different cadherins. While engineered cell systems provide excellent experimental tools to observe pattern formation in cell populations, computational models may be leveraged to more systematically explore the key parameters that drive the emergence of different patterns. We developed and validated two- and three-dimensional agent-based models (ABMs) of spheroid patterning for cells engineered with a bidirectional signaling circuit that regulates N- and P-cadherin expression. The model was used to predict how varying initial cell seedings, cadherin induction probabilities, or homotypic adhesion strengths between cells impact spheroid patterning, and unsupervised machine learning techniques were used to map system parameters to unique spheroid patterns. The model was then deployed to design new synthetic cell signaling circuits based on a desired final multicell pattern.

## INTRODUCTION

Multicell patterning processes in embryonic development and tumorigenesis are regulated by dynamic cell-to-cell signaling and differential adhesion between cell subpopulations (1, 2). Steinberg and colleagues (3) previously proposed the differential adhesion hypothesis to explain how patterns emerge at the multicell level in heterogenous mixtures of cells. Specifically, they proposed that cell subpopulations self-organize to minimize the adhesive free energy between cells such that more adhesive cell subpopulations are surrounded by less adhesive ones (3–5). The differential adhesion hypothesis has been validated experimentally (3–7) and incorporated into computational models of cell-sorting and morphogenesis (8–11). However, this conceptual model does not account for how dynamic cell-to-cell signaling processes, which regulate adhesion protein expression, may yield emergent patterns that differ from those predicted by purely equilibrium considerations (i.e., minimization of adhesive free energy).

The ability for a cell to adhere to another cell may be time- and space-dependent based on cell-to-cell signaling that alters cell expression of adhesion proteins. In embryonic development, for example, cell-to-cell interactions initiate transcriptional processes that regulate expression of specific adhesion proteins, leading to the segregation of cell subpopulations into organ-specific tissue layers (12–19). Similarly, in many cancers, signaling cascades cause individual cells to downregulate expression of the cell adhesion protein E-cadherin, leading to epithelial-mesenchymal transition (EMT) (20–24). EMT creates the opportunity for differential adhesion between epithelial cells and the mesenchymal counterparts they can generate, impacting the spatial configuration of these subpopulations throughout the tumor (25–28). Emergent cell patterning within tumors can impact tumor growth, migration, and response to treatment (20, 28–34). In these natural biological systems, the multicell patterns that form may depend on the initial ratio of cell subpopulations and/or the order of signaling events in the system, but more work is needed to explore such potential dependencies.

Synthetic biology approaches have been used to impart self-organizing properties to multicellular systems (2, 35–41). These systems provide an ideal setting to explore pattern formation resulting from dynamic cell-to-cell signaling because the cell interactions are clearly defined and programmable. A recent example of interest is the bidirectional signaling system engineered by Toda et al. (2), in which sender cells drive expression of a particular cadherin in receiver cells that then reciprocally drive expression of a different cadherin in the initial sender cells (Figure 1A). The authors observed that the relative homotypic and heterotypic cadherin interaction strengths directly impacted observed spheroid patterns. When the network was engineered with N-cadherin (Ncad) and P-cadherin (Pcad), which both have high homotypic and low heterotypic adhesion strengths, a “core/pole” pattern arose. Specifically, Ncad-expressing receiver cells formed a core decorated by smaller clusters, or poles, of Pcad-expressing sender cells and surrounding non-adhesive sender cells (2). When the system was instead engineered with high and low expression of E-cadherin (Ecad), creating cells with high (but differential) homotypic and heterotypic adhesion, a “core/shell” pattern arose. Specifically, high-Ecad receiver cells formed a core encapsulated by a shell of low-Ecad sender cells (2). In both systems, the number of poles and core/shell spheroids formed varied across trials with the same initial conditions (2), but the factors that led to this variable outcome were not systematically investigated.

**Figure 1.**
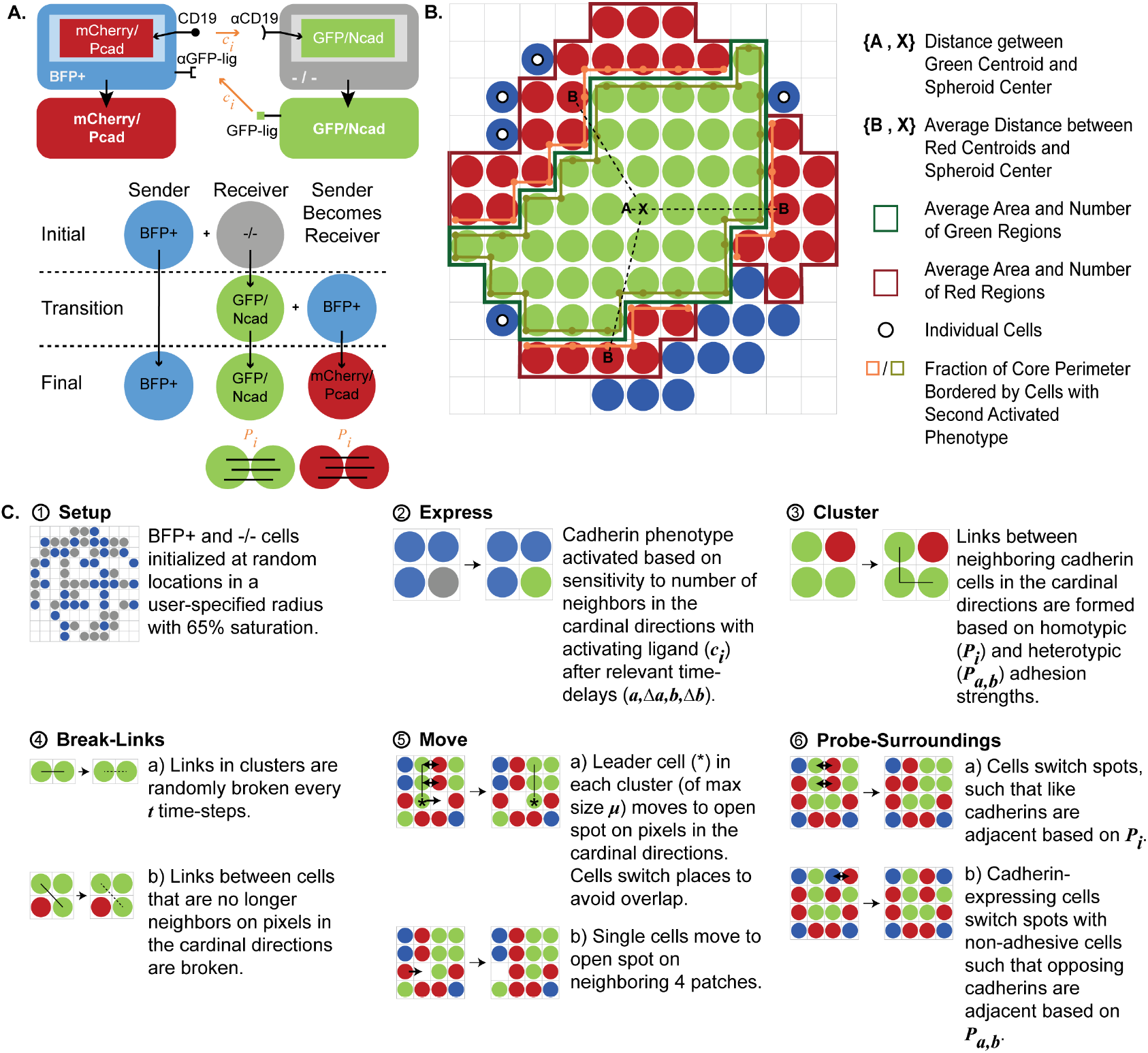
Synthetic biology functionality can be encoded in an agent-based model of cell-to-cell signaling within a spheroid. **(A)** A schematic of the synthetic bidirectional signaling circuit engineered by Toda et al. (2) and implemented in the agent-based model (ABM) is shown. Spheroids are seeded with non-adhesive “sender” BFP+ cells and “receiver” −/− cells (labeled −/− to indicate no fluorescent protein expression). Sender BFP+ cells can induce GFP/Ncad expression in −/− receiver cells, and newly formed GFP/Ncad cells behave as sender cells to reciprocally induce mCherry/Pcad expression in BFP+ cells (at which point BFP+ cells behave as receiver cells). The final spheroid composition can include GFP/Ncad, mCherry/Pcad, or BFP+ cells (that never contacted a GFP/Ncad cell and thus failed to transition to the mCherry/Pcad phenotype). In this ruleset, the model cadherin induction constant (*c*_*i*_) controls the probability of GFP/Ncad induction in −/− cells based on neighboring BFP+ cells and the probability of mCherry/Pcad induction in BFP+ cells based on neighboring GFP/Ncad cells. The model homotypic adhesion strength (*P*_*i*_) controls the probability of forming junctions between cells expressing like cadherins on adjacent grid spaces. **(B)** Metrics used to analyze ABM outputs are represented schematically. Regions of a given color comprising three or more cells are defined as cell clusters. **(C)** The rules executed at each ABM timestep are depicted. Parameters, indicated in bold italics, are defined in Table 1.

**Table 1.** ABM parameter values.

| Parameter | Value | Definition | Justification for Value |
| --- | --- | --- | --- |
| $a$ | 10 ticks | Delay for GFP/Ncad phenotype to be induced in the system. | Estimated from Toda et al. (2018); set such that GFP/Ncad cells start to appear at ~26 ticks in the model. |
| $\Delta a$ | 10 ticks | Time to express Ncad. | |
| $b$ | 25 ticks | Delay for mCherry/Pcad to be induced in the system. | Estimated from Toda et al. (2018); set such that mCherry/Pcad cells start to appear at ~42 ticks in the model. |
| $\Delta b$ | 10 ticks | Time to express Pcad. | |
| $\mu$ | 5 cells | Maximum movable cluster size (in number of cells). | Manually set such that smaller homotypic clusters can move throughout the spheroid to improve likelihood of homotypic clusters merging to form larger homotypic clusters. |
| $t$ | 3 ticks | Delay allotted after links are randomly broken in clusters to be reformed. | Manually set such that sufficient time is allowed for individual cell agents to move after a link is broken, so that new clusters can be formed in the spheroid. |
| $c_i$ | 0.20 (unitless) | Induction constant in probability of expression formula that controls the probability of a cadherin phenotype being induced in a receiver cell based on the number of neighboring sender cells. $c_a$ and $c_b$ are used to refer to induction constants for GFP/Ncad and mCherry/Pcad phenotypes, respectively, in figures 6-10. | Identified by the evolutionary algorithm. |
| $P_i$ | 0.95 (unitless) | Probability of homotypic adhesion in induced cadherin cell types. $P_a$ and $P_b$ are used to refer to individual homotypic adhesion strengths of GFP/Ncad and mCherry/Pcad, respectively, in Figures 6-10. | |
| $P_{a,b}$ | 0.05 (unitless) | Probability of heterotypic adhesion between GFP/Ncad and mCherry/Pcad cells. | Manually set to allow few heterotypic links to be formed between GFP/Ncad and mCherry/Pcad cells. |

While it would be impossible to exhaustively explore the effects of perturbing different system parameters experimentally, computational models provide an efficient method for predicting unique multicellular patterns for different parameter combinations (42–46). ABMs are a particularly useful computational approach for modeling heterogeneous cell types and spheroid patterning because ABMs can represent individual cells within a population and predict how cell type-specific behaviors produce spatial patterns at the multicell level (8–10, 28, 47–54). Here, we developed two-dimensional (2D) and three-dimensional (3D) ABMs of the Ncad/Pcad bidirectional signaling system engineered by Toda et al. (2). Cells were modeled as expressing different cadherins with a specified time delay in response to heterotypic receptor-ligand interactions and as differentially adhering to like or unlike cells based on the homotypic and heterotypic adhesion properties of the cadherins. Model parameters were tuned to fit experimental observations we made for different seeding densities of initial cell types. Unsupervised machine learning was then employed to map different combinations of perturbed model parameters (e.g., characterizing cadherin induction and adhesion strengths) to specific spheroid patterns predicted by the ABM, including the previously described core/pole and core/shell patterns and others resembling a soccer ball, bull’s eye, or stripes. The ABM was also used to design synthetic cell-to-cell signaling circuits that encode specified multicellular patterns, demonstrating a potential practical application of this modeling approach.

## RESULTS

### 2D and 3D ABMs predict the impacts of dynamically activated differential adhesion on spheroid patterning

2D and 3D ABMs were developed based on the synthetic Pcad/Ncad circuit engineered by Toda et al. (2) (Figure 1A). The models were seeded with “sender” and “receiver” cells. Sender cells expressed blue fluorescent protein (BFP+), CD19 ligand, and green fluorescent protein (GFP) receptor. Receiver cells expressed the CD19 receptor (−/−). In response to CD19 receptor-ligand binding between the two cell types, receiver −/− cells express Ncad and GFP to become GFP/Ncad cells. GFP/Ncad cells then behave as sender cells and reciprocally promote Pcad and mCherry expression in BFP+ cells (that then behave as receiver cells) via GFP ligand-receptor binding, forming mCherry/Pcad cells. Hence, in this synthetically engineered multicell system, both cell types can function as either sender or receiver cells based on receptor-ligand binding interactions with adjacent cells, and the terms “sender” and “receiver” are used to clarify a cell’s role during specific steps in bidirectional signaling. Final spheroid patterns were quantified using metrics, such as total fraction of spheroid area occupied by each cell type, average areas of homotypic cell clusters, and average distances between homotypic cell clusters and the spheroid center (Figure 1B). Homotypic cell “clusters” were defined as more than two cells of the same cell type adjoined by cell-to-cell junctions. The “core” cell type of a spheroid was defined as the cell type of the homotypic cluster with minimum distance to all other homotypic clusters in the spheroid.

Interactions among cells in the ABM were encoded as six rules incorporating nine model parameters that execute on each timestep of the simulation (Figure 1C, Table 1). Cadherin phenotypes were induced in the model based on induction constants (*c*_*i*_) that control the probability of cadherin induction in a receiver cell based on the number of neighboring sender cells. Over the time course of the model, neighboring cadherin-expressing cells form junctions with each other based on homotypic (*P*_*i*_) and heterotypic (*P*_*a,b*_) adhesion strengths encoded as probabilities of adhesion in the model. To recapitulate the Ncad/Pcad spheroids observed in Toda et al. (2), we assumed that both cadherin cell types have an equal induction constant (*c*_*i*_) and equal homotypic adhesion strength (*P*_*i*_). In later sections, we explore how spheroid patterning is impacted by assigning unequal values of *c*_*i*_ and *P*_*i*_ to the GFP/Ncad phenotype (*c*_*a*_, *P*_*a*_) and to the mCherry/Pcad phenotype (*c*_*b*_, *P*_*b*_). All simulations that were compared to *in vitro* experiments were run for 100 timesteps, where one timestep represents 30 minutes, because *in vitro* spheroids were observed for 50 hours. All other simulations were run until no changes were observed in the spheroid pattern, and the specific model runtime for each set of simulations is denoted in corresponding figure captions.

### Published data and an evolutionary algorithm were used to parameterize the ABM

Parameters in the 2D ABM were either estimated based on published experimental data in Toda et al. (2) or computationally tuned with an evolutionary algorithm (EA) (Table 1). The time delay for cadherin phenotypes to be induced (*a* for GFP/Ncad and *b* for mCherry/Pcad) and the time to express each cadherin (Δ*a*, Δ*b*) were manually adjusted in the 2D ABM according to published data in Toda et al. (2). Specifically, these parameters were set such that in simulations of spheroids seeded with 100 BFP+ and 100 −/− cells, GFP/Ncad cells formed after 13 hours and mCherry/Pcad cells developed adjacent to GFP/Ncad cells after 21 hours, the respective times reported by Toda et al. (2). Because the remaining five parameters (*c*_*i*_, *P*_*i*_, *P*_*a,b*_, *μ*, *t*) could not be determined from experimental data, these parameters were first manually approximated in the 2D ABM as values yielding outputs qualitatively similar to published images of spheroids from the Ncad/Pcad bidirectional signaling system in Toda et al. (2) (i.e., characterized by a GFP/Ncad central core flanked by mCherry/Pcad clusters and individual BFP+ cells). Then, a univariate parameter sensitivity analysis was implemented for the 2D ABM to determine which of these parameters had the largest relative impact on spheroid patterning. Each of the five parameters was increased or decreased by 10% from its base value, and the effects of these perturbations on the numbers and average areas of green and red clusters were determined (Figure 2A-D). This analysis identified the cadherin induction constant (*c*_*i*_) and the homotypic adhesion strength of cadherin cell types (*P*_*i*_) as the two parameters whose variation produced the largest changes in all four measured outputs.

**Figure 2.**
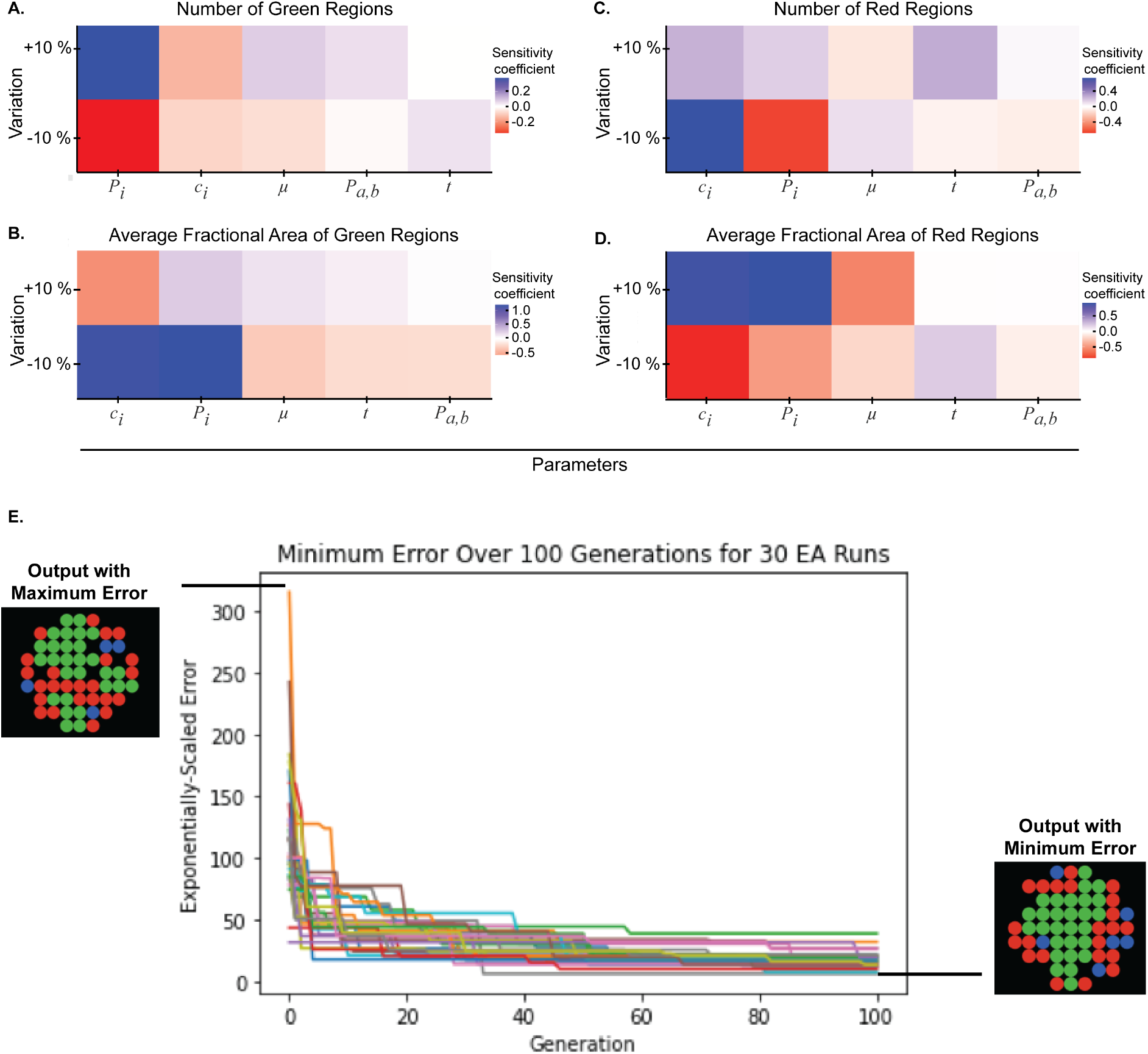
Parameter sensitivity analysis and optimization improves model fit to *in vitro* data. Parameters that could not be estimated based on experimental data (*c*_*i*_, *P*_*i*_, *P*_*a,b*_, *μ*, *t*) were perturbed by a factor 0.9 and 1.1 and the sensitivity coefficient was quantified for the following outputs: **(A)** number of green regions, **(B)** average fractional area of green regions, **(C)** the number of red regions, **(D)** and the average fractional area of red regions. Sensitivity coefficients are reported as the average of 100 model runs, each run for 100 timesteps. **(E)** An evolutionary algorithm (EA) was used to tune the parameters *c*_*i*_ and *P*_*i*_ by minimizing the error between model outputs and experimental images of 200-cell spheroids seeded with a 1:1 ratio of BFP+ and −/− cells. The EA was run 30 times for a population of 20 model combinations over 100 generations. Within each generation, the error for each parameter combination was averaged over 10 runs for 100 timesteps. Sample model graphical outputs generated from the parameter combinations with maximum or minimum error are shown.

Values for the parameters *c*_*i*_ and *P*_*i*_ were determined using an evolutionary algorithm (EA) that optimized the 2D ABM prediction of patterns that were experimentally observed in 200-cell spheroids seeded with a 1:1 ratio of BFP+ and −/− cells. A detailed description of the EA implementation is provided in *Materials and Methods*. A plot of 30 EA runs over 100 generations demonstrates that the EA accomplished a substantial reduction in error (difference between model prediction and data) to identify values for *c*_*i*_ and *P*_*i*_ (Figure 2E). The parameter values optimized for the 2D ABM were also used in the 3D ABM.

### ABMs predict spheroid patterns across a range of different initial conditions that match *in vitro* results

2D and 3D ABMs were used to predict spheroid patterns when the BFP+ : −/− cell seeding ratio was varied (1:9, 1:4, 1:1, 4:1, 9:1) for a fixed total number of seeded cells (60 or 200 cells) (Figure 3A). For the 1:1 ratio with 200 cells, the model predicted a GFP/Ncad core surrounded by mCherry/Pcad poles and individual BFP+ cells. For the 1:1 ratio with 60 cells, the model predicted two neighboring poles, one of GFP/Ncad cells and one of mCherry/Pcad cells, with surrounding individual BFP+ cells. In spheroids with a minority of seeded BFP+ cells, fewer mCherry/Pcad cells were generated, leading to fewer and smaller mCherry poles. Conversely, in spheroids with a majority of seeded BFP+ cells, fewer GFP/Ncad cells were generated, leading to smaller GFP/Ncad clusters that were typically located farther from the spheroid center. Model predictions aligned well with observations from *in vitro* experiments performed using the cell lines engineered by Toda et al. (2). Indeed, the average fractional blue, red, and green areas of the *in vitro* spheroids and the model predictions were statistically similar to each other for the 1:9, 1:1, and 9:1 ratios (Figure 3B, 3C, 3D). Moreover, 2D and 3D model predictions were statistically similar, suggesting that the 2D model sufficiently represented the synthetic cell signaling system.

**Figure 3.**
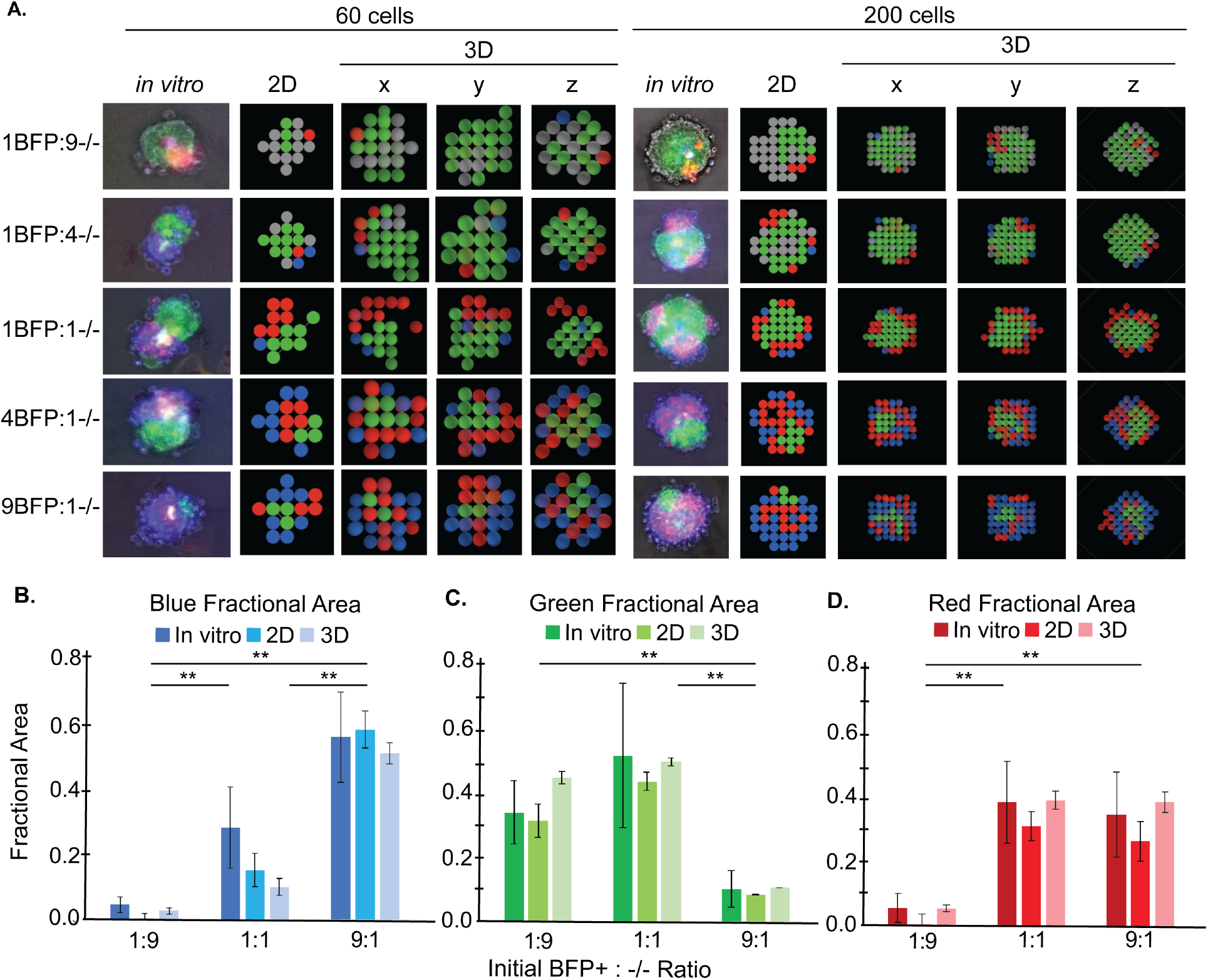
ABM qualitatively and quantitatively recapitulates *in vitro* pattern formation. **(A)** Comparisons are shown of representative cross-sectional images of *in vitro* spheroids with 2D or 3D ABM outputs. 3D ABM *x*-, *y*-, and *z*-cross-sections provide 2D views of the spheroid cores. *In vitro* cross-sectional images are microscopy images of a single focal plane, without deconvolution, in the approximate center of the spheroid. Comparisons of **(B)** blue, **(C)** green, and **(D)** red fractional areas between *in vitro*, 2D ABM, and 3D ABM images for varying initial BFP+:−/− seeding ratios (1:9, 1:1, 9:1) and 200 cells. ABM-predicted fractional areas for the 2D and 3D models were calculated as an average of 100 model runs, each run for 100 timesteps, for each experimental condition. Fractional areas from *in vitro* images were calculated from a sample of 9, 10, and 10 images for the 1:9, 1:1, and 9:1 ratios, respectively. A one-way Kruskal-Wallis test followed by multiple-comparisons testing was used to compare fractional areas between all *in vitro*, 2D, and 3D ratios (** indicates *p* < 0.001). Error bars indicate within-group standard deviation.

Comparison of fractional areas of induced phenotypes among the 1:9, 1:1, and 9:1 ratios indicates that a small number of sender cells is required to induce cadherin phenotypes in the majority of receiver cells. For example, despite there being fewer BFP+ cells in the 1:9 ratio, the total GFP/Ncad fractional area was not significantly different from that of the 1:1 ratio (Figure 3C). Similarly, despite the total fractional area of GFP/Ncad cells being significantly lower in the 9:1 ratio than in the 1:1 ratio (Figure 3C), the total mCherry/Pcad fractional area was not significantly different from the 1:1 ratio (Figure 3D). These results suggest that sender BFP+ cells in the 1:9 ratio and sender GFP/Ncad cells in the 9:1 ratio disperse sufficiently over the model’s time course such that sender BFP+ and sender GFP/Ncad cells experience enough interactions with receiver cells that the total fractional area of phenotypes induced by these sender cells is similar to that in the 1:1 ratio. This hypothesis was tested by quantifying the positions and movements of GFP/Ncad cells over time in the 2D ABM for 1:1, 4:1, and 9:1 BFP+ : −/− seeding ratios. For ratios with BFP+ cells in the majority (4:1 and 9:1), GFP/Ncad regions were significantly farther from the spheroid center than for the 1:1 ratio and, thus, more dispersed throughout the spheroid (Figure 4A). Additionally, after the GFP/Ncad phenotype had been induced in the majority of −/− cells (∼13 hours or 26 timesteps in the model), the fraction of GFP/Ncad cells that moved per timestep in the 9:1 ratio was larger than that in the 4:1 and 1:1 ratios (Figure 4B). This result suggests that simulated cell clusters composed of fewer cadherin-expressing cells were more mobile due to fewer movement-restricting intercellular interactions. Increased mobility permits more interactions of GFP/Ncad cells with BFP+ cells, ultimately producing as many mCherry/Pcad cells for the 9:1 condition as for the 1:1 condition.

**Figure 4.**
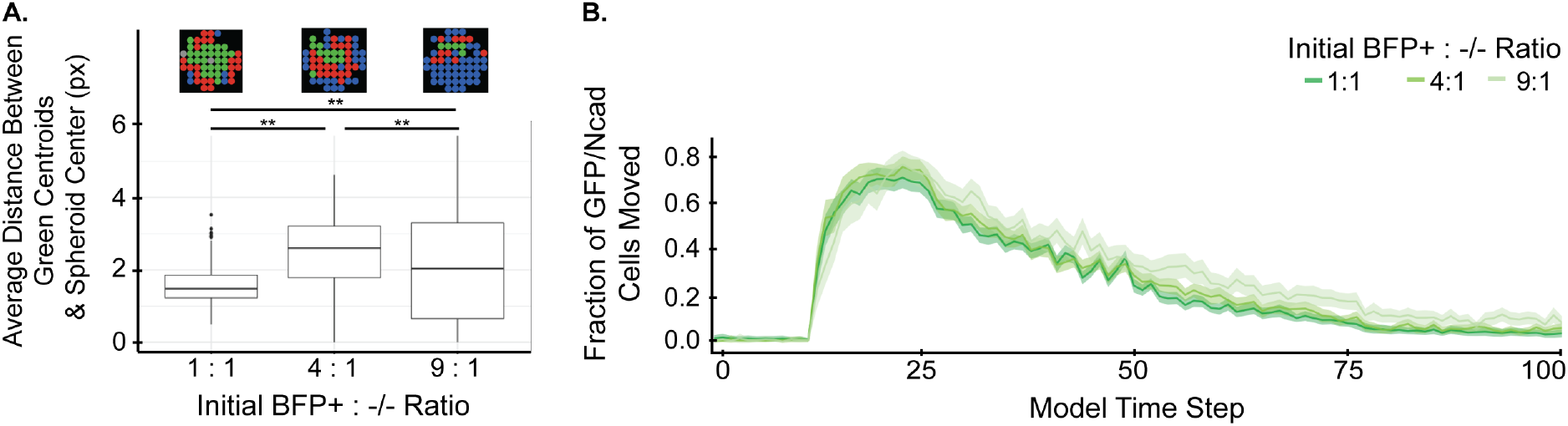
The relative fraction of seeded BFP+ sender cells impacts the size and mobility of GFP/Ncad clusters. **(A)** The average distance between green region centroids and spheroid centers was quantified for increasing ratios of initial BFP+ seeded cells in 200-cell spheroids. Data are shown as a boxplot distribution for 100 model runs, each run for 100 timesteps. The groups were compared using a one-way Kruskal-Wallis test followed by individual Dunn’s tests for posthoc testing (** indicates *p* < 0.001). **(B)** From timestep 26 onwards, the number of green cells that move per tick is generally higher in spheroids with a 9:1 ratio of initial BFP+ cells to −/− cells. Metrics were calculated as the average from 100 model runs, each for 100 timesteps. Ribbons in the line plot represent the 95% confidence interval calculated using the standard error of the mean.

The model also predicted that the total number of seeded cells affected final spheroid patterns (Figure 5A). As the total number of cells in the spheroid increased, the number of green regions formed also increased (Figure 5B). Simultaneously, the average fractional area of green regions in the spheroid decreased, and the average fractional distance of green regions to the center of the spheroid increased (Figure 5C and Figure 5D). These trends suggest that as the overall cell number increases, the central GFP/Ncad core partitions into smaller pieces that become interspersed throughout the spheroid. This result is further explored in Figure 7.

**Figure 5.**
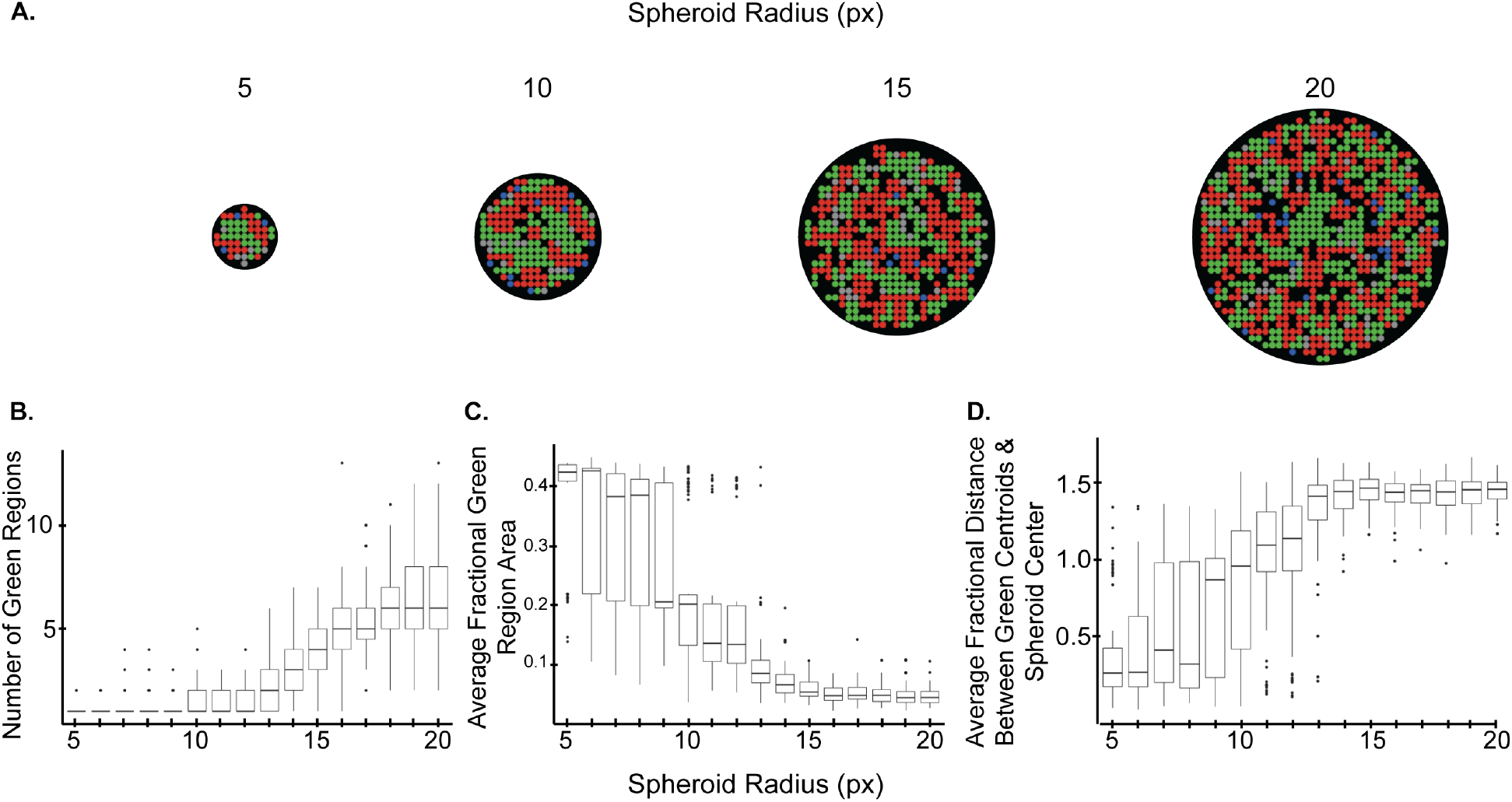
Increasing the radius of spheroids with a 1:1 BFP+:−/− seeding ratio leads to deviation from the core/pole configuration. The **(A)** number of green regions, **(B)** average fractional area of green regions, and **(C)** average fractional distance between green region centroids and spheroid center were quantified as a function of model spheroid radius (reported in units of pixels, px). Data are shown as boxplots of values for 100 model runs, each run for 100 timesteps.

### Dynamic cell-to-cell signaling and homotypic adhesion strength impact pattern formation

To test the hypothesis that the dynamic expression of different cadherin isoforms impacts spheroid patterning, three different signaling circuits were explored (Figure 6A). In circuit 1, the cadherins were constitutively expressed, as opposed to expression via induction by juxtracrine signaling. In circuit 2, only unidirectional signaling was present such that seeded mCherry/Pcad sender cells induced the GFP/Ncad phenotype in seeded −/− receiver cells. In circuit 3, the full bidirectional signaling circuit was modeled, with seeded BFP+ sender cells inducing GFP/Ncad expression in seeded −/− receiver cells and newly formed GFP/Ncad cells inducing mCherry/Pcad expression in BFP+ cells. In this circuit, the induction constants for GFP/Ncad and mCherry/Pcad phenotypes were parametrized separately (*c*_*a*_ and *c*_*b*_, respectively, instead of *c*_*i*_). In the simulations discussed for circuit 3 in this section, *c*_*a*_ = *c*_*b*_ = *c*_*i*_, but *c*_*a*_ and *c*_*b*_ are varied in later *Results* sections. For each circuit, three homotypic adhesion strength profiles were tested, in which the homotypic adhesion strengths of GFP/Ncad and mCherry/Pcad phenotypes were parametrized separately (*P*_*a*_ and *P*_*b*_, respectively, instead of *P*_*i*_). In profile A, GFP/Ncad and mCherry/Pcad cell types exhibited maximal homotypic adhesion and minimal heterotypic adhesion (*P*_*a*_ = *P*_*b*_ = 1.0, *P*_*a,b*_ = 0.0), giving rise to core/pole spheroids, in which one adhesive cell type assembled in a core and the other assembled in smaller peripheral clusters. In profile B, GFP/Ncad cells exhibited maximal homotypic adhesion, mCherry/Pcad cells exhibited minimal homotypic adhesion, and heterotypic adhesion was maximal (*P*_*a*_ = 1.0, *P*_*b*_ = 0.0, *P*_*a,b*_ = 1.0). In profile C, GFP/Ncad cells exhibited minimal homotypic adhesion, mCherry/Pcad cells exhibited maximal homotypic adhesion, and heterotypic adhesion was maximal (*P*_*a*_ = 0.0, *P*_*b*_ = 1.0, *P*_*a,b*_ = 1.0). Both profiles B and C gave rise to core/shell spheroids, in which the more adhesive cell type formed a core surrounded by a complete shell of the less adhesive cells (Figure 6A). Combinations of each signaling circuit and adhesion profile yielded 9 rulesets. For all rulesets, simulations were seeded with a 1:1 ratio of initial cell types for a fixed total of 200 cells.

**Figure 6.**
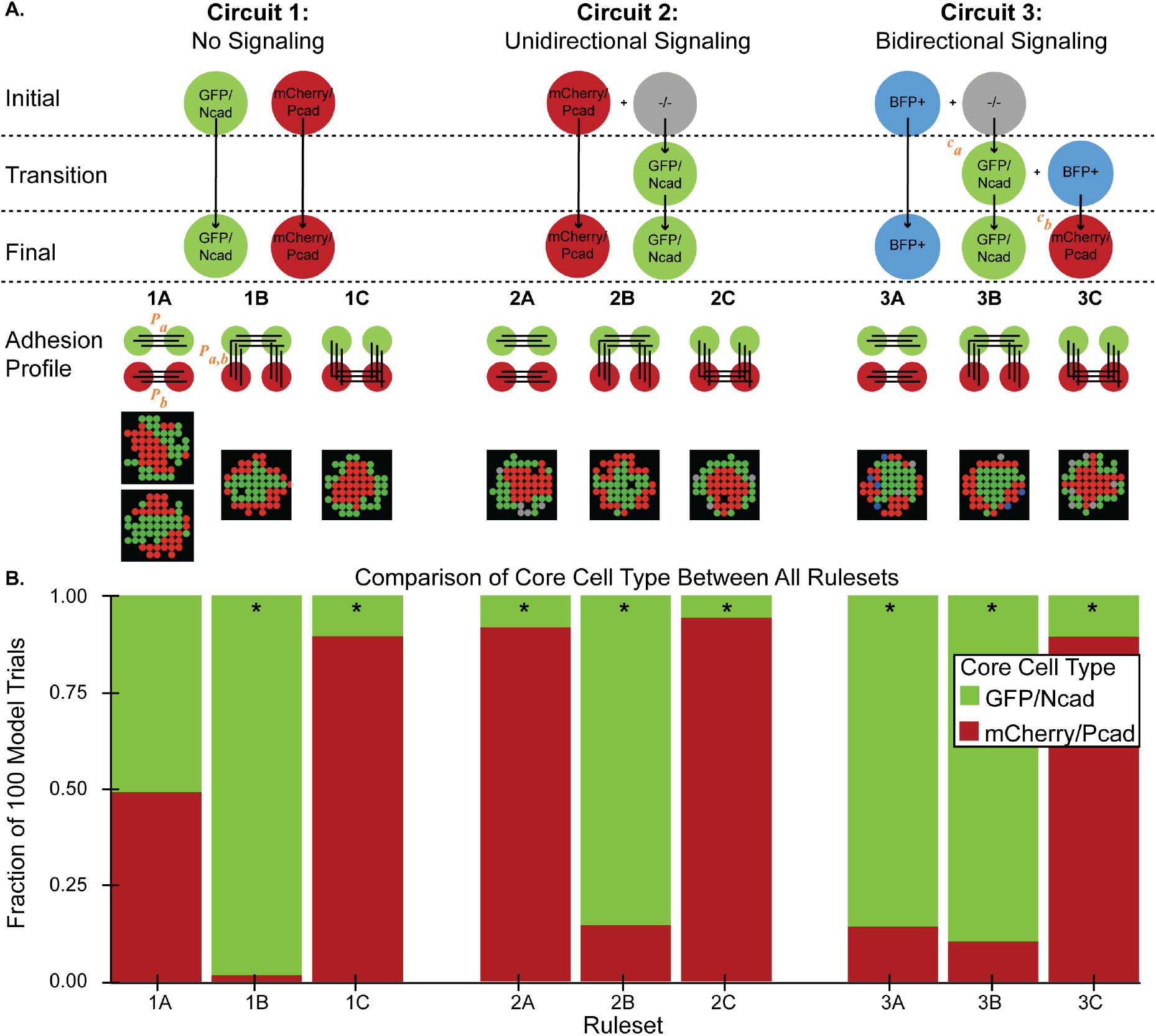
Dynamic cell-to-cell signaling has differential effects on core/pole and core/shell spheroids. **(A)** A schematic of three different ABM signaling circuits and three adhesion profiles is shown. Circuit 1 contains constitutive cadherin expression and no juxtracrine signaling. Circuit 2 includes unidirectional signaling. Circuit 3 includes bidirectional signaling (as described in Figure 1A). In Circuit 3, induction constants for GFP/Ncad and mCherry/Pcad phenotypes are parametrized separately (*c*_*a*_ and *c*_*b*_, respectively). In each adhesion profile, the homotypic adhesion strengths of GFP/Ncad and mCherry/Pcad phenotypes are also parametrized separately (*P*_*a*_ and *P*_*b*_, respectively). All depicted adhesion profiles have a strength of 1.0. Adhesion profile A gives rise to core/pole spheroids, whereas adhesion profiles B and C give rise to core/shell spheroids. The combination of circuit type and adhesion profile leads to nine rulesets (1A, 1B, 1C, 2A, 2B, 2C, 3A, 3B, and 3C). **(B)** The model was run 100 times, each for 100 timesteps, for each of the nine rulesets. Each ruleset was seeded with 200 cells at a 1:1 ratio of the seeding cell types. The fraction of trials with either GFP/Ncad or mCherry/Pcad spheroid core was compared between rulesets using a Chi-Squared test with df = 8; significant differences between observed and expected fraction (0.50) of core cell types were found across all nine rulesets with *p* < 0.001. The expected fraction was set to 0.5 because the null hypothesis states that signaling has no effect on which cell type forms the spheroid core. * indicates groups with residuals > 1.

The effects of uni- and bi-directional signaling between cells were explored by comparing the core cell types in spheroids for circuits 1-3 (Figure 6B). Adhesion profile A tended to produce core/pole spheroids for which the core cell type varied based on the signaling circuit. Ruleset 1A gave rise to either GFP/Ncad or mCherry/Pcad cores, whereas in Rulesets 2A and 3A, the adhesive phenotype that was first present in the circuit formed the spheroid core. Thus, the adhesive cell type that is present longer in the spheroid exhibited a higher likelihood of condensing into a central core. The same trends for core cell type were not observed for adhesion profiles B and C, which tended to produce core/shell patterns. In core/shell spheroids, the more adhesive phenotype formed the spheroid core in > 90% of model simulations regardless of the signaling circuit (Figure 6B). Thus, for profiles B and C, the signaling dynamics produced outcomes that were consistent with the differential adhesion hypothesis in 200-cell spheroids seeded with a 1:1 ratio.

### Varying cell seeding in Ruleset 3B leads to patterns that violate the differential adhesion hypothesis

To test whether it is possible to generate spheroids that violate the differential adhesion hypothesis, the 2D ABM was used to predict patterns of cell subpopulations when seeding ratios and total cell numbers in Ruleset 3B were varied. Ruleset 3B was chosen because it is based on the fundamental premise of the differential adhesion hypothesis, in which two phenotypes with differing homotypic adhesion strengths self-organize. As the relative amount of seeded −/− cells and spheroid radius increased, the likelihood that adhesive cells would enclose non-adhesive cells increased (Figure 7A). In the 1:1 ratio group, regardless of spheroid radius, a small number of simulated spheroids (∼12-20%) contained one group of non-adhesive red cells enveloped by adhesive green cells, while virtually no spheroids contained non-adhesive −/−, or gray, cells enveloped by adhesive green cells. For the 1:4 and 1:9 ratios, the fraction of spheroids with groups of non-adhesive red cells enclosed by adhesive green cells decreased significantly, and the overall fraction of spheroids with groups of non-adhesive gray cells enclosed by adhesive green cells increased significantly, with the latter effect becoming more apparent as the spheroid radius increased. This trend is partially explained by the presence of fewer seeded BFP+ cells, which leads to fewer −/− cells transitioning to the GFP/Ncad phenotype and mCherry/Pcad phenotypes. Due to the increased tendency of smaller green clusters to spontaneously arise at the periphery of the spheroid, non-adhesive cells surrounding each individual green cluster appeared to be trapped in the center of the spheroid, giving rise to situations where non-adhesive cells appeared enclosed by adhesive cells. This hypothesis is further reinforced by the fact that when the induction constant of the GFP/Ncad phenotype was increased (*c*_*a*_), a single, larger green core formed and the spheroid patterns resembled those observed in the 1:1 group (Figure 7B, 7C). When the induction constant of the GFP/Ncad phenotype (*c*_*a*_) was larger, fewer BFP+ sender cells were required to induce the GFP/Ncad phenotype in −/− receiver cells, and a larger central GFP/Ncad core formed more quickly in the spheroid, instead of the slower formation of GFP/Ncad clusters and their subsequent enclosure of non-adhesive cells.

**Figure 7.**
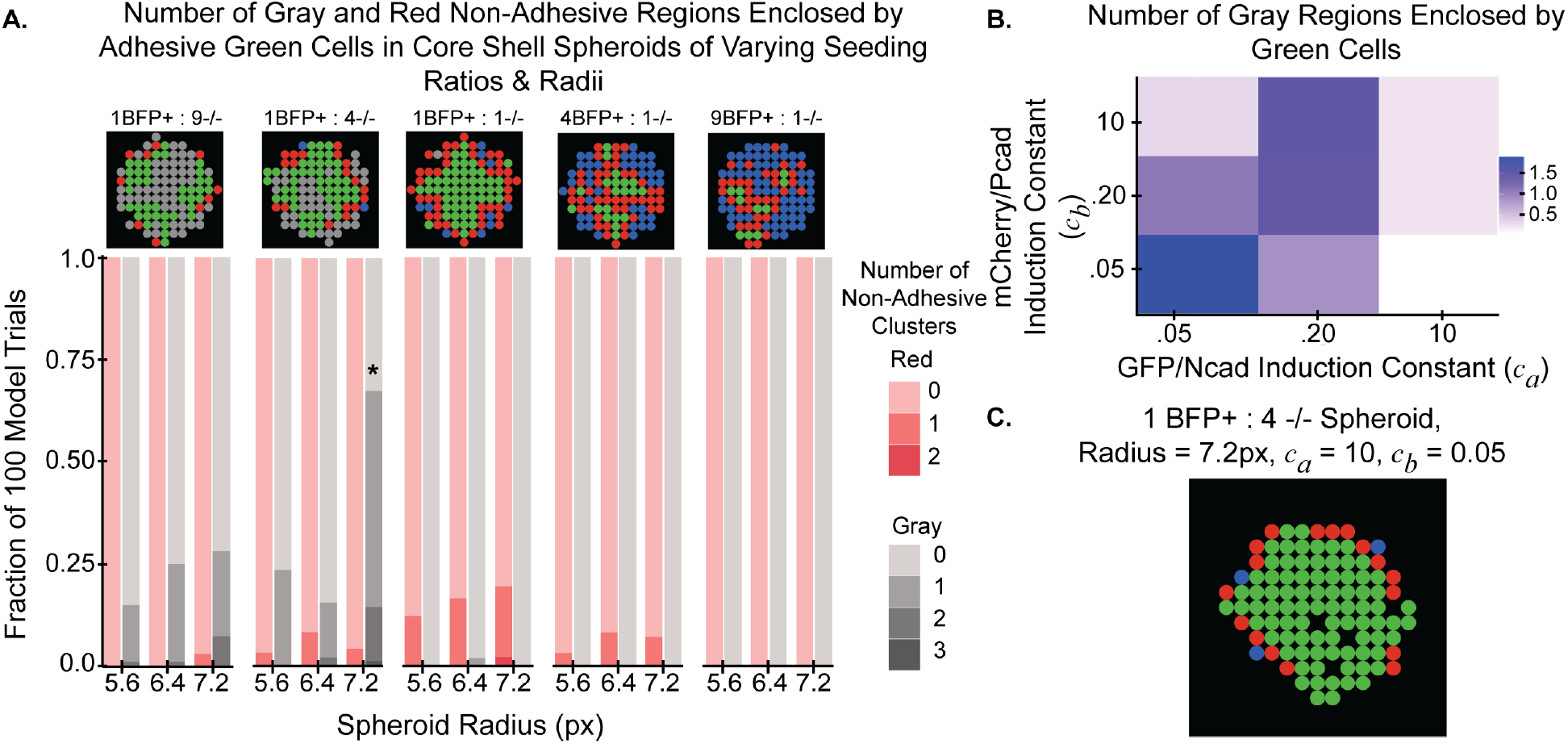
Increasing the relative fraction of seeded −/− cells in Ruleset 3B leads to instances where non-adhesive cell types are surrounded by adhesive cell types. **(A)** The number of gray and red non-adhesive cell clusters with greater than 75% enclosure by green adhesive cells was averaged for 100 model runs, each run for 200 timesteps, for spheroids with radii 5.6, 6.4, and 7.2 pixels (px) and varying initial BFP+:−/− ratios (1:9, 1:4, 1:1, 4:1, 9:1). All groups were compared with a Chi-Squared test with df = 14, resulting in a *p* < 0.001 (* indicates groups with greater than 10% contribution to the chi statistic. **(B)** *c*_*a*_ and *c*_*b*_ were simultaneously varied over values of 0.05, 0.2, and 10 in spheroids with radius 7.2 px and initial BFP+:−/− ratio of 1:4. Total number of gray clusters with greater than 75% enclosure by green cells was quantified for 100 model runs, each run for 200 timesteps, for each model condition. **(C)** A representative output of a simulated spheroid with radius 7.2 px, initial BFP+:−/− seeding ratio of 1:4, and *c*_*a*_ = 10 and *c*_*b*_ = 0.05.

### Homotypic adhesion strengths impact heterogeneity in spheroid patterns

Systematically covarying the homotypic adhesion strengths of GFP/Ncad and mCherry/Pcad cells (*P*_*a*_ and *P*_*b*_, respectively) and setting *P*_*a,b*_ = 0 in the 2D ABM for signaling circuit 3 (bidirectional signaling) produced a range of different spheroid patterns. To display the results of these calculations, heatmaps were created to represent the numbers and average areas of green or red regions in the spheroids (Figure 8A). Recall that the extremes of these maps describe core/shell (*P*_*a*_ = 1.0 and *P*_*b*_ = 0.0 or *P*_*a*_ = 0.0 and *P*_*b*_ = 1.0) and core/pole patterns (*P*_*a*_ = *P*_*b*_ = 1.0). With *P*_*a*_ > *P*_*b*_, there were typically fewer and larger green regions than red regions, but the average number and area of green regions was not as low as those observed in the core/pole pattern with a red core. Conversely, with *P*_*b*_ > *P*_*a*_, there were fewer and larger red regions than green regions, but the average number and area of red regions was not as low as those observed in the core/pole pattern with a green core. Deviations from core/pole and core/shell patterns could occur when *P*_*a*_ and *P*_*b*_ have non-extreme values.

**Figure 8.**
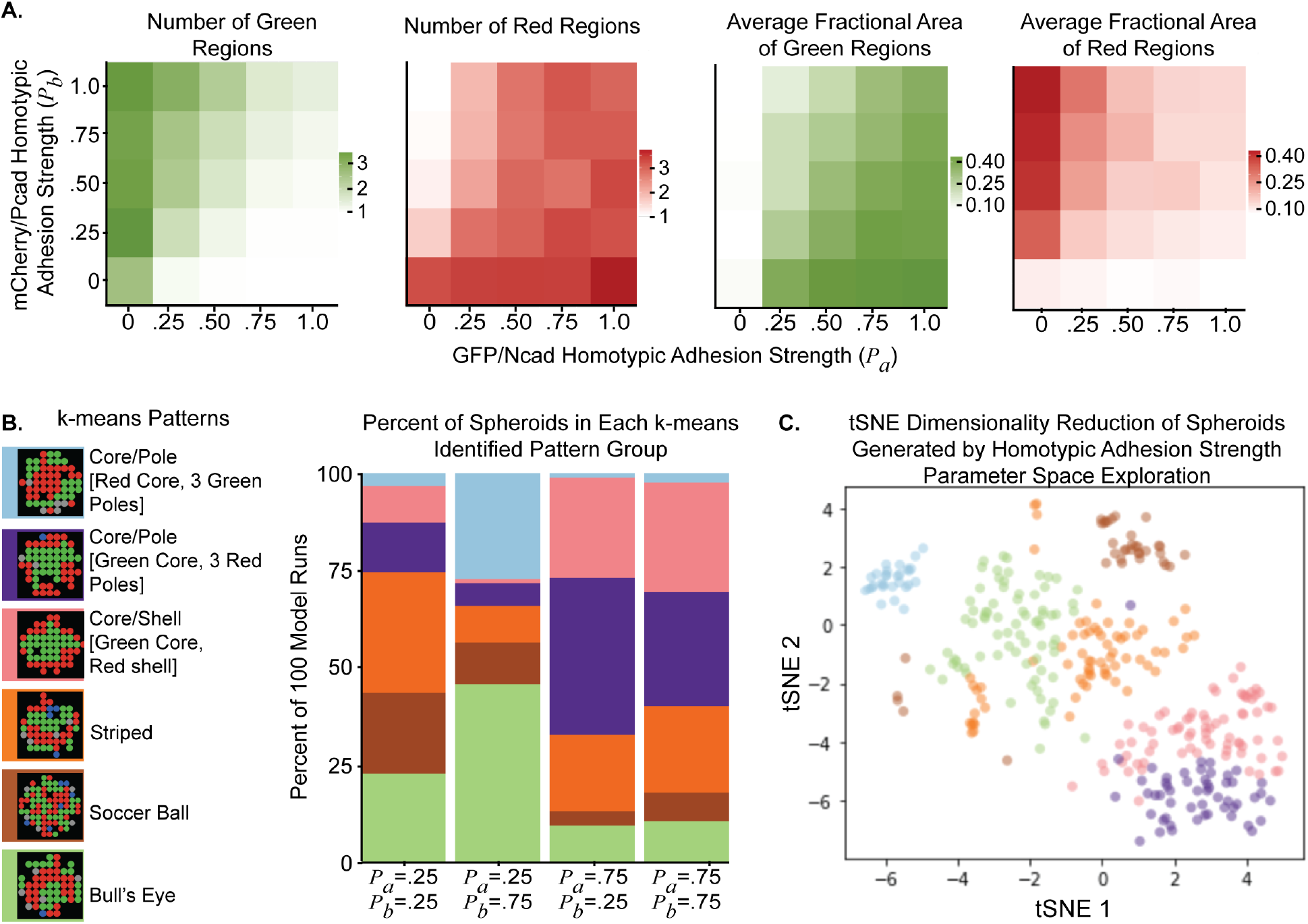
Covarying homotypic adhesion strengths of activated cell types in Ruleset 3A leads to a variety of different patterns. **(A)** Homotypic adhesion strengths of the first and second activated phenotypes (*P*_*a*_, *P*_*b*_) were simultaneously varied with values of 0.0, 0.25, 0.50, 0.75, and 1.0 for signaling circuit 3 (bidirectional signaling circuit) in the 2D ABM. For each parameter combination, the simulations were run for 200 cells at a 1:1 BFP+:−/− ratio, and were run 100 times, each for 100 timesteps. The average values of the indicated metrics were plotted in heat maps. **(B)** Results from this parameter space exploration were clustered using a k-means algorithm into 6 group, with representative ABM outputs shown for each group and color coded as the key for this graph. **(C)** A t-distributed stochastic neighbor embedding (tSNE) algorithm was used to show where the k-means clusters projected in two dimensions. The tSNE algorithm was run with the perplexity parameter = 100.

To identify the different spheroid patterns formed, we performed k-means clustering, an unsupervised machine learning technique, to identify six groups within a subset of the data, namely the samples with [*P*_*a*_, *P*_*b*_] = [0.25, 0.25], [0.25, 0.75], [0.75, 0.25], and [0.75, 0.75]) (Figure 8B). Each of the six groups corresponded to a distinct spheroid pattern: a core/pole pattern with a red core and three green poles, a core/pole pattern with a green core and three red poles, a core/shell pattern with green core and red shell, a striped pattern of alternating green and red linear regions, a soccer ball pattern of circular green and red regions, and a bull’s eye pattern with radially alternating green and red shells. As an alternative way to visualize the k-means clustering results, the data were projected in two dimensions using t-distributed stochastic neighbor embedding (tSNE), with pattern identities retained from k-means clustering (Figure 8C). These results also visually support the binning of spheroids into more than just two categories.

Returning to the results shown in Figure 8B, we note that each combination of homotypic adhesion strengths generated simulation outputs that exhibited all six patterns identified by the clustering algorithm described above. The frequency of each pattern, however, depended on the parameters. When the GFP/Ncad phenotype had relatively weak homotypic adhesion (e.g., [0.25, 0.25] and [0.25, 0.75]), striped, soccer ball, and bull’s eye patterns emerged more frequently than in conditions when the GFP/Ncad phenotype had stronger homotypic adhesion (e.g., [0.75, 0.25] and [0.75, 0.75]). When the GFP/Ncad phenotype had relatively weak homotypic adhesion, the frequency of heterotypic interactions between cells was greater, allowing for a higher frequency of patterns characterized by longer borders between heterotypic cells.

In conditions where differential adhesion was present (e.g., [0.25, 0.75] and [0.75, 0.25]), the most frequent patterns involved the more adhesive cells being surrounded by the less adhesive cells. With *P*_*a*_ = 0.75 and *P*_*b*_ = 0.25, the most frequent patterns were the core/pole and core/shell spheroids with a green core. With *P*_*b*_ = 0.75 and *P*_*a*_ = 0.25, the most frequent patterns were the core/pole spheroids with a red core and the bull’s eye spheroids with a red core. This trend suggests that when differential adhesion is present, but the adhesive strength of the second induced phenotype is not maximal, hybrids of the core/shell pattern arise.

When both cell types had relatively weak homotypic adhesions (e.g., [0.25, 0.25]), the patterns were relatively evenly distributed, indicating that when less homotypic adhesion is present in the system, the increased opportunity for heterotypic interactions leads to greater heterogeneity in pattern formation. Alternatively, when both cell types had relatively strong homotypic adhesion (e.g., [0.75, 0.75]), patterns with core/pole structures occurred more frequently, indicating a skew towards maximal contact between like cells.

### Cadherin induction constants impact core/pole and core/shell patterns

Varying the induction constants of GFP/Ncad and mCherry/Pcad cells (*c*_*a*_ and *c*_*b*_, respectively) was also predicted to impact the final numbers of green and red regions and their average areas (Figure 9A). For Ruleset 3A (bidirectional signaling core/pole ruleset) and when *c*_*a*_ = 0.05 and *c*_*b*_ = 10, one green region with a relatively small area was formed, and one red region with a larger area was formed. When *c*_*b*_ was only slightly larger than *c*_*a*_, multiple small green regions were formed, and one or two red regions were formed, suggesting the possibility that different patterns may occur for different parameter combinations. Seven groups were identified by k-means clustering (Figure 9B). The first two patterns included a green core and three or four red poles. The third pattern comprised spheroids with a green core and red shell. The fourth pattern exhibited an inversion of the first two, with a red core and two or more green poles. The fifth pattern included one red core and one green pole. The sixth included a red core and single green and gray peripheral cells. Finally, the seventh pattern included striped or soccer ball patterns. As in Figure 9A, when *c*_*a*_ << *c*_*b*_, a red core formed with one or multiple green poles. However, when *c*_*a*_ was sufficiently high, core/pole structures with a green core and multiple red poles formed more frequently. To produce an alternative visualization, tSNE was used to reduce the dimensionality of the system (Figure 9C). tSNE results further supported the existence of multiple subtypes of spheroids. Modifying the cadherin induction constants affects the overall spheroid pattern because various combinations of *c*_*a*_ and *c*_*b*_ directly impact the final ratio of BFP+, −/−, GFP/Ncad, and mCherry/Pcad cells in the spheroid (Figure 9D).

**Figure 9.**
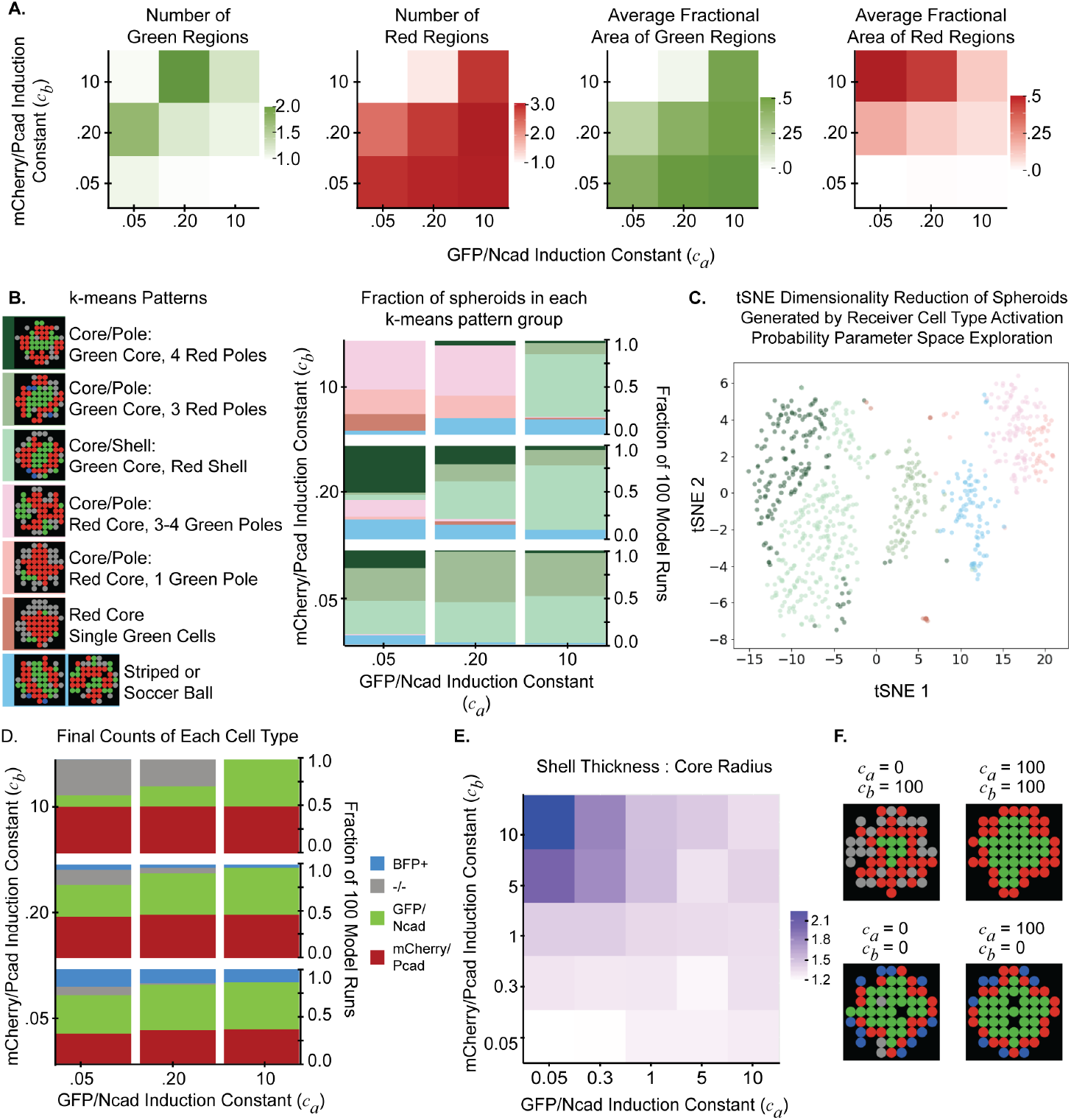
Cadherin induction constants (*c*_*i*_) control the number of poles in core/pole spheroids and the ratios of shell thickness to core radius in core/shell spheroids. **(A)** The cadherin induction constants (*c*_*a*_, *c*_*b*_) were simultaneously varied across values of 0.05, 0.2, and 10 for Ruleset 3A (bidirectional signaling core/pole ruleset) in the 2D ABM. For each parameter combination, simulations were seeded with 200 cells at a 1:1 BFP+:−/− ratio and run 100 times, each for 100 timesteps. The average from 100 model runs of the following metrics were plotted for each parameter combination: number of green regions, number of red regions, average fractional area of green regions, and average area of red regions. **(B)** Results from this parameter space exploration were clustered using a k-means algorithm into 7 groups, with representative ABM outputs shown for each group and color coded as the key for this graph. **(C)** t-distributed stochastic neighbor embedding (tSNE) algorithm was used to project the data into two dimensions with colors for spheroid cluster types retained from the k-means clustering analysis in panel (B). The tSNE algorithm was run with perplexity = 100. **(D)** The average fraction of BFP+, −/−, GFP/Ncad, and mCherry/Pcad cells in spheroids were calculated for each parameter combination. (**E**) The cadherin induction constants (*c*_*a*_, *c*_*b*_) were simultaneously varied across values of 0.05, 0.3, 1, 5, and 10 for Ruleset 3B (bidirectional signaling core/shell ruleset) in the 2D ABM. For each parameter combination, simulations were seeded with 200 cells at a 1:1 BFP+:−/− ratio and run 100 times, each for 100 timesteps. **(F)** Representative images of spheroids from extremes of 9E are shown.

The parameters *c*_*a*_ and *c*_*b*_ were also varied for Ruleset 3B (bidirectional signaling core/shell ruleset). The ratio of the shell’s thickness to the core’s radius progressively increased as *c*_*b*_ became larger than *c*_*a*_ (Figure 9E and 9F), which is consistent with more mCherry/Pcad cells at the final time point. Because this parameter describes sensitivity of a cell to its neighbors, and the number of neighbors in 3D is larger than in 2D, results from these 2D model calculations may not translate directly to a 3D model.

### Adjusting ABM parameters enables customization of multicell patterns in spheroids

To demonstrate the utility of the model for developing design principles for spheroid engineering, we used the 2D ABM to design multicellular spheroids with desired patterns. To control the overall spheroid configuration, the homotypic adhesion strengths of induced cell types should be controlled (Figure 10A). The ABM predicted that, to produce a core/pole pattern, both adhesion molecules should have homotypic adhesion strengths > 0.75. In order to produce a core/shell pattern, one induced phenotype must have homotypic adhesion > 0.75, while the other induced phenotype must have a homotypic adhesion strength < 0.25. To produce soccer ball, striped, or bull’s eye patterns, both activated phenotypes must have homotypic adhesion strengths in the range 0.25-0.75.

**Figure 10.**
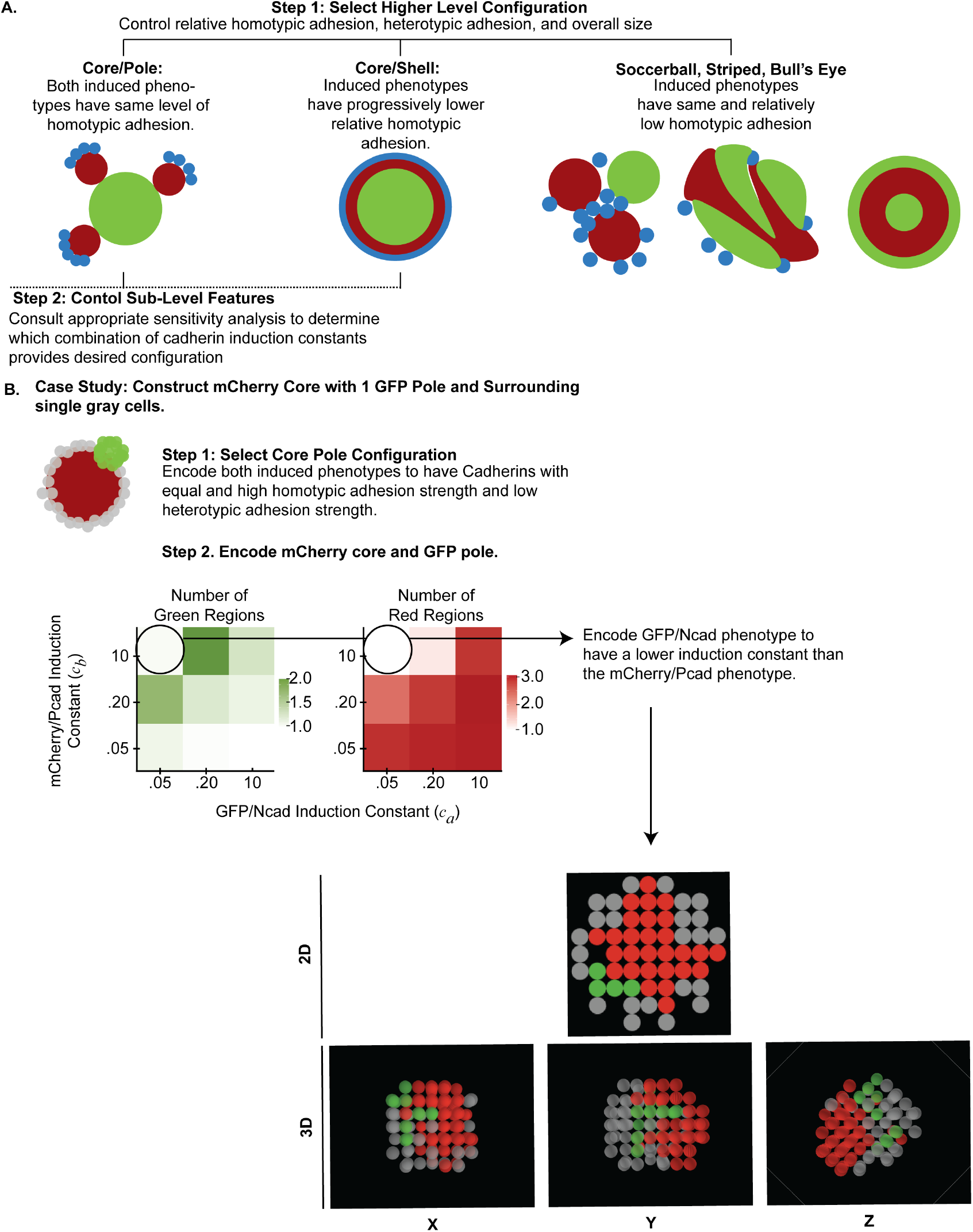
ABMs may be used to design synthetic signaling circuits encoding customizable core/pole, core/shell, soccer ball, striped, and bull’s eye spheroid patterns. **(A)** For a desired spheroid pattern (e.g., core/pole, core/shell, soccer ball, striped, or bull’s eye), homotypic and heterotypic adhesion strengths of cell types can be modulated. Features of the core/pole and core/shell configurations, such as the number of poles and ratio of shell thickness to core radius, may be further controlled by modifying the induction constants of cadherin phenotypes in the system. **(B)** An outline of the process to create a structure with mCherry/Pcad core, GFP/Ncad pole, and surrounding single cells is shown.

The results of Figure 10B demonstrate how bivariate parameter explorations of the 2D ABM can guide the forward design of a user-specified spheroid with an mCherry/Pcad core, one GFP/Ncad pole, and surrounding −/− cells. Based on the results of the model, the user should set *P*_*a*_ and *P*_*b*_ to be maximal and equal to achieve a core/pole configuration and engineer the system such that *c*_*a*_ << *c*_*b*_ to achieve an mCherry core and one GFP/Ncad pole.

## DISCUSSION

We created an ABM of bidirectional cell-to-cell interactions that drive patterning of multicellular spheroids inspired by the signaling circuits engineered by Toda et al. (2). The ABM provides an alternative to time-consuming cell culture experimentation by predicting patterns that can arise from a multicell signaling system as a result of adjusting initial conditions, signaling interactions, and adhesion strengths alone and in combination. We used the model to explore the impact of dynamic cell-to-cell signaling on adhesion-driven pattern formation, and we observed the emergence of new patterns that were inconsistent with the classical differential adhesion hypothesis. Moreover, we used machine learning to map combinations of homotypic adhesion strengths to unique spheroid patterns. Additionally, we found that modulating the induction constants for the GFP/Ncad and mCherry/Pcad phenotypes could control the number of poles in core/pole patterns or the ratio of shell thickness to core radius in core/shell patterns. Our study supports the use of computational models to identify parameters that will yield customized spheroid patterns. Because these parameters have physical meaning, they may be used as design constraints for multicell, synthetic circuits. For example, homotypic and heterotypic adhesion strengths may be controlled by selecting specific cadherin isoforms (6). Similarly, promoter regions for synthetic transgenes may be selected according to the intended phenotype induction constants (55).

Previous studies have leveraged computational models to explore cell patterning in heterogeneous systems (10, 43–45, 47–49, 56, 57). The differential adhesion hypothesis has been computationally modelled using a Cellular Potts Model framework (8, 9). These models have demonstrated how cell subpopulations with varying levels of cohesiveness self-assemble, and how environmental variables such as temperature, shear force, and pressure affect these self-assembly processes. CompuCell3D, for example, is a multi-scale model based on this Cellular Potts Model framework that recapitulates morphogenesis in a wide array of tissues and species, and considers unique organ developments as a result of cell-to-cell signaling (10). Taylor et al. (11) developed an ABM to explore boundary formation and border sharpening between subtypes in the Eph-ephrin signaling system, which is critical in embryogenesis. Their model presents a range of configurations that arise between Eph- and ephrin cells, and as a function of varying the homotypic and heterotypic adhesion strengths between cell types. Although these published models have thoroughly explored pattern formation in heterogeneous systems with static cell phenotypes, they have not considered how patterns may be affected by the modulation of differential adhesion via known, programmable cell-to-cell signaling interactions. Our model differs from prior efforts because it simulates phenotypic transitions controlled by a synthetic gene circuit, which allowed us to explore how the phenotype induction probabilities, timing, and sequencing of dynamic cell-to-cell differential adhesion drives pattern formation.

By simulating a multicell system containing genetically engineered signaling circuits that control cell-to-cell adhesion, we demonstrated the utility of agent-based modeling in helping to design such circuits to achieve desired multicell patterns. Depending on the desired pattern (e.g., core/shell, core/pole), different combinations of adhesion molecules or methods to induce their expression may be selected. Duguay et al. (6) and Foty & Steinberg (7) quantified the relative homotypic adhesion and heterotypic adhesion strengths of cadherin molecules, and these studies may be used as reference for selecting adhesion molecules for synthetic circuit designs. In order to control details of the pattern, such as the number of poles and ratio of the shell thickness to the core radius, the induction constant of cells may be adjusted. Matsuda et al. (55) defined a parameter similar to ours as the “promoter coefficient” in their synthetic cell-cell system. This coefficient describes the probability of gene expression upon cell-to-cell contact with the sender cell. Cellular features related to this model parameter may include the copy number of the promoter of interest in different cell lines and the likelihood of methylation based on the insertion region.

Aspects of the basic modeling approach utilized here will also be useful for the study of endogenous cell signaling circuits that regulate cell-to-cell adhesion. A specific application of relevance in cancer biology is that of epithelial-mesenchymal transition (EMT), which aberrantly occurs in numerous carcinomas (e.g. breast cancer, pancreatic ductal adenocarcinoma) and is thought to promote metastasis and chemoresistance (23, 24). During EMT, epithelial-derived cancer cells lose polarity and expression of cellular adhesion molecules to become more mesenchymal. This process is highly regulated by signaling and transcriptional networks that can be initiated by growth factors, cell-matrix interactions, and tumor microenvironmental factors (e.g., low oxygen tension). Thus, ABMs capable of incorporating the signaling dynamics that lead to EMT and the resultant effects on differential cell-to-cell adhesion could be used to better understand the morphogenesis of primary tumors and to potentially understand the processes that lead tumors of different compositions to shed metastases (28, 54). Our modeling framework would extend previous relevant computational models of how differential adhesion impacts tumor metastasis and cell migration (58).

There are limitations of both our computational model and in the *in vitro* system. The computational model does not simulate the effects of cell proliferation nor the fact that induced cadherin expression could decrease after loss of contact between sender and receiver cells. The ABM also assumes that cadherin expression is uniform among cells, but realistic heterogeneity of expression within a population of even clonal cells could impact pattern formation. These issues could be addressed in future model implementations. For example, the distributions of expression of different adhesion molecules among cells could be quantified by flow cytometry or single-cell RNA sequencing, and this data could inform individual simulated cell behaviors in the ABM. Experimental limitations, including an inability to count the individual cells within spheroids, also limited the degree to which we could quantitatively compare model results with experimental data. Furthermore, our model and the experimental spheroids do not contain diffusible morphogen cues, which are instructive in the patterning of natural (non-synthetic) multicell biological systems (41). Although this is a limiting factor for the applicability of the ABM developed here to the study of naturally-occurring systems, future extensions of the ABM could incorporate diffusible morphogen cues for inducing cell differentiation (44, 45). An additional limitation of our model is that signaling dynamics were encoded as simple time delays for cadherin expression after cell contacts were made. In the future, it will be worthwhile to incorporate more realistic models of intracellular signaling, as has been done in recently published multiscale ABMs that integrate information across intracellular and intercellular scales (59–62). The need to solve differential equations describing reaction kinetics in each cell within the ABM will greatly increase the computational resources needed to simulate the system as well as the number of model parameters. These are widespread issues encountered in multiscale, mechanistic biological models (42, 58), and the tradeoffs must be carefully weighed in designing the structure of such models. Encouragingly, recent studies have presented novel methods for reducing the computational burden of complex mechanistic models by using neural networks and hybrid continuum-based modeling approaches (63–66).

An emerging trend in the field of multiscale computational modeling of biological systems is the integration of ABMs with machine learning and other data science approaches to more richly exploit high throughput experimental and simulated data. We demonstrated the utility of coupling an ABM with machine learning through our use of k-means clustering, which mapped cell-specific parameters in the ABM (homotypic adhesion strengths and cadherin induction constants) to different tissue-level spheroid patterns. Previous studies have similarly used ABMs in conjunction with machine learning methods to study and predict multicellular interactions (46, 67). Future iterations of our ABM could combine data-driven modeling and machine learning to further inform rule development (63, 64), model calibration (68), and parameter exploration (67) in order to create an even more comprehensive and predictive model.

Our study motivates several follow-up questions about multicell patterning that can be pursued by extending and redeploying our ABM. We can further explore the mechanisms by which other multicell patterns that have been observed in nature arise (e.g., stripes, soccer balls, and bull’s eyes). Moreover, it would be interesting to explore how varying initial distributions of differentially activated cell types impact the final spheroid pattern formed. Additionally, the introduction of environmental variables, such as diffusible chemokines that control chemotaxis, could be incorporated into the ABM for additional complexity and biological relevance. Finally, the model could be applied to simulate natural systems such as collections of tumor cells displaying various degrees of epithelial-mesenchymal transition.

## MATERIALS AND METHODS

### Cell lines and propagation

L929 murine adipocytes engineered with a bidirectional cell-to-cell signaling circuit based on the synthetic notch (synNotch) receptor system (2) were generously provided by Dr. Wendell Lim (University of California, San Francisco). Cells were maintained in DMEM (ThermoFisher) supplemented with 10% fetal bovine serum (VWR), 1 mM L-Glutamine, 100 units/mL penicillin, and 100 μg/mL streptomycin. Cell lines were used within 25 passages and maintained in a 37°C, 5% carbon dioxide ThermoForma i160 incubator.

### Spheroid culture

A total of 60 or 200 cells (BFP+:−/− ratios = 1:9, 1:4, 1:1, 4:1, 9:1) were seeded per well in Ultra Low Attachment 96-well microplates (Corning™ #7007, Corning, N.Y). Cells aggregated into spheroids at the bottom of the well. 12 spheroids were plated per condition. For 200-cell spheroids, 50 μL of 4 cells/μL cell suspension and 50 μL of DMEM were loaded in each well. For 60-cell spheroids, 50 μL of 1.2 cells/μL cell suspension and 50 μL of DMEM were loaded in each well.

Spheroids were imaged 50 hours after plating using a Zeiss AxioObserver Z1 widefield microscope at 10× magnification. Six to seven representative spheroids were selected for imaging for each experimental condition, by selecting wells without plastic particles from plate manufacture and wells in which all three cell colors were easily detected due to spheroid orientation. For each spheroid, a phase contrast image and three fluorescence images were taken: BFP (DAPI filter set), GFP (AF488 filter set), and mCherry (AF546 filter set). Zen Zeiss software was used to merge phase, BFP, GFP, and mCherry images.

### ABM design and implementation

2D and 3D on-lattice ABMs were developed to simulate collections of cells containing the synthetic signaling circuit represented in Figure 1. An ABM platform was chosen because this modeling technique allows for prediction of spheroid-level patterns based on individual cell-to-cell interactions. The 2D and 3D models were developed using NetLogo software (69).

The ABMs treat cells as individual agents in a spheroid. The 2D ABM simulates the cells interacting in a plane through the central axis of the spheroid, whereas the 3D ABM simulates all cells throughout the spheroid. Each grid space in the model represents a 10 μm × 10 μm area capable of accommodating one cell. Each model timestep represents 30 minutes such that 100 timesteps represent the 50-hour period over which spheroids were studied *in vitro*. Unless otherwise noted, simulations were run for 100 timesteps because this is the time over which spheroids were observed *in vitro*. Seeded model agents represent BFP+ sender cells and −/− receiver cells, which can become mCherry/Pcad and GFP/Ncad cells, respectively. State transitions were encoded by setting Boolean values of private variables. Intercellular interactions were encoded as six rules, characterized by ten parameters, that execute on each timestep (Figure 1C, Table 1).

Figure 1C describes the rules of the ABM. At the initial timestep, the “setup” method randomly places BFP+ and −/− cells within a user-specified radius and according to a user-specified ratio and percent saturation (percent of cell-occupied grid spaces). On every subsequent timestep of the model, cells move to the open grid space (of eight neighboring spaces in 2D and 26 neighboring spaces in 3D) closest to the center of the simulation space to represent the gravitational and adhesive forces that tend to pull them together at the bottom of a low-attachment well. If there are no open spaces available on the neighboring eight grid spaces of a cell, then that cell will not move.

Clusters of cells containing fewer than a threshold number of cells (*μ*) identify the cell closest to the spheroid center as a cell that moves to a randomly-selected space on one of its neighboring eight grid spaces (or 26 grid spaces in 3D) that is unoccupied. If there are no open spaces surrounding the leader cell, the cluster will not move. Like cells in a contiguous cluster follow the leader as a connected unit, displacing cells in previously occupied grid spaces as needed to avoid two cells occupying the same grid-space simultaneously. Displaced cells move randomly to the positions vacated by the cluster that moved in response to a leader cell move. As the model is on-lattice, movement is discrete (i.e., between grid spaces), and each grid space is occupied by only one cell at a given time.

After a time delay (*a*), −/− receiver cells can convert to GFP/Ncad cells depending on the number of neighboring BFP+ sender cells. Similarly, after another time delay (*b*), BFP+ sender cells can express mCherry/Pcad depending on the number of neighboring sender GFP/Ncad cells. Neighboring cells are defined as cells occupying the eight surrounding grid spaces in 2D or the 26 surrounding grid spaces in 3D. The probability of phenotypic induction (*P*_*induction*_) for receiver cells in contact with sender cells was determined as

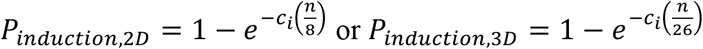

where *c*_*i*_ is a tunable parameter that controls the probability of cadherin expression induction, and *n* is the number of neighbors with the activating phenotype. If the cadherin phenotype is induced, cells change colors and express Ncad or Pcad after specified time delays (Δ*a* and Δ*b*, respectively). In this baseline ruleset, both cadherin phenotypes have the same induction constant, *c*_*i*_.

Cadherin-expressing cells attach to other cadherin-expressing cells based on homotypic and heterotypic adhesion probabilities (*P*_*i*_ and *P*_*a,b*_, respectively). These parameters dictate the probability that a cadherin-expressing cell will form a link with a neighboring like or unlike cadherin-expressing cell within a single simulation timestep, or tick. Additionally, when a cell expressing one type of cadherin is wedged between two cells expressing another type of cadherin, the cells switch places based on the homotypic adhesion probability such that the cells with the two like cadherins occupy adjacent grid spaces. Similarly, if a cell expressing one type of cadherin is separated from a cell expressing a different cadherin by a non-cadherin expressing cell, cells switch places based on the probability *P*_*a,b*_ such that cells expressing the two different cadherins occupy adjacent spaces. This type of position switching behavior has been experimentally observed (4). In the baseline ruleset, both cadherin phenotypes have the same homotypic adhesion strength, *P*_*i*_. Additionally, to provide the opportunity for cells to leave one cluster and join other clusters, 50% of the cadherin interactions across a grouping of connected cells were randomly broken at each timestep, and cadherin interactions between groups of cells could only be reformed after a specific number of ticks specified by the parameter *t*.

### *In vitro* and *in silico* quantification of multicell patterns

A variety of characteristics that describe the spatial patterns of cells in the *in vitro* spheroids and *in silico* spheroids were quantified (Figure 1B). A MATLAB pipeline was developed to quantify the following characteristics for each color/channel from images of *in vitro* spheroids: fractional area, total area, and distance between region centroid and spheroid center. In order to do this, thresholds were applied to images to isolate purely green, red, or blue pixels. Then, pixel areas and centroids of contiguous regions for each color were computed based on the MATLAB method ‘RegionProps,’ which computes these metrics using a connected components algorithm.

The same features were quantified in the 2D ABM simulations. Similarly, the MATLAB connected components ‘RegionProps’ algorithm was used to quantify the total fractional area, average area of homotypic cell clusters, and average distance between the centroid of homotypic clusters and the spheroid center for each color channel. A cluster was defined as a group of adjacent cells containing more than two cells. Additionally, the number of non-clustered cells in each color channel and total cell count of each color channel was quantified. Finally, a contiguous area ratio, which represents the fraction of the border of a spheroid core that is surrounded by cells expressing the non-core forming cadherin, was also computed. In the 3D ABM, the total fractional volume of each color channel was quantified. This metric was computed as the cell count of a certain color channel divided by the total number of cells in the spheroid.

### Parameterization of the ABM using experimental data

Table 1 summarizes the ABM parameter values and how they were determined. The parameters *a*, *b*, Δ*a*, and Δ*b* were estimated according to data published in Toda et al. (2), which indicated that for 200-cell spheroids seeded with an initial 1:1 BFP+:−/− ratio, GFP/Ncad cells appeared after 13 hours and mCherry/Pcad cells appeared adjacent to GFP/Ncad cells after 21 hours (2). Values for *a*, Δ*a*, *b*, and Δ*b* were determined such that the model matched these results at the corresponding time points of 26 ticks and 42 ticks, since one model tick (or timestep) represents 30 minutes. The remaining five model parameters (*c*_*i*_, *P*_*i*_, *P*_*a,b*_, *μ*, and *t*) could not be determined from experimental data. As a first approximation, these parameters were manually adjusted to baseline values so that the 2D ABM yielded spheroids that were qualitatively similar to images of the Ncad/Pcad signaling circuit published in Toda et al. (2). Specifically, values for *μ* and *t* were manually set to encourage continual movement of homotypic clusters in the spheroid over the 100-tick period and prevent the model from reaching a “static” state composed of smaller homotypic clusters at early timepoints.

### Univariate parameter sensitivity analysis

A univariate sensitivity analysis was performed to identify which of the five parameters that could not be experimentally determined (*c*_*i*_, *P*_*i*_, *P*_*a,b*_, *μ*, and *t*) most control model outputs. Each model parameter was perturbed, one at a time, by a factor of 0.9 or 1.1, and a sensitivity coefficient (*S*) was calculated as

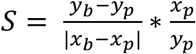

where *x*_*b*_ and *x*_*p*_ represent baseline and perturbed parameter values, respectively, and *y*_*b*_ and *y*_*p*_ represent the baseline and perturbed values of the model output, respectively. Sensitivity coefficients were computed for the following model outputs: number of green regions, average fractional area of green regions, number of red regions, and average fractional area of red regions. The mean sensitivity for each model output was computed from 100 model runs and reported for the 0.9 and 1.1 perturbation of each of the 10 parameters.

### Computational tuning with an evolutionary algorithm

An evolutionary algorithm (EA) was used to tune parameters that could not be determined from published data. Due to the high complexity and computational expense of parameter tuning, only the top two sensitive parameters from the univariate sensitivity analysis (*c*_*i*_ and *P*_*i*_) were selected for tuning. The EA finds a parameter combination that minimizes an error function through a natural selection process wherein optimal parameter combinations in parent generations are selected for cross-over to create offspring parameter combinations in subsequent generations. The error, *E*, that the EA minimized was calculated as

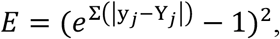

where *y*_*j*_ represents the value of a calculated model output, and *y*_*j*_ represents the experimental value for that output. The following model outputs (*j*) were used in the error equation: total fractional green area, total fractional red area, total fractional blue area, average fractional green region area, average fractional red region area, number of green regions, and number of red regions. The exponential form of the error equation was used to encourage EA convergence. The EA was run for a population size of 20 (i.e., 20 randomly generated sets of initial guesses for *c*_*i*_ and *P*_*i*_), with 10 parallel ABM runs to compute an average error and 100 generations for each set of initial guesses. At each generation, a cross-over rate of 50% and mutation rate of 80% was used. This process (20 sets of initial guesses × 10 ABM runs × 100 generations) was repeated 30 times, and the combination of *c*_*i*_ and *P*_*i*_ producing the overall lowest error was chosen to model the Ncad/Pcad signaling circuit developed by Toda et al. (2). This approach was used to account for stochasticity of the ABM and EA and to reduce the likelihood of the EA becoming stuck in local error minima. We developed our own Python script for the EA and used the NL4Py Python library (70) to enable communication between Python and NetLogo. To improve run time, our EA implementation ran the ABM in parallel across multiple computing units.

### Comparison of ABM predictions with experimental data

The 2D and 3D ABMs were validated against experimental data by simulating different BFP+ : −/− seeding ratios (1:9, 1:4, 1:1, 4:1, 9:1) for 60 and 200 total cells, replicating the conditions used for *in vitro* experiments. Simulations for each set of conditions were run 100 times, and the average results from these runs were analyzed. The locations of each simulated cell were saved at the end of each model run, and metrics were quantified as described above. Fractional areas/volumes of each color were compared between *in vitro* results and ABM predictions.

### Rule set implementations

Different rulesets were explored using the 2D ABM by varying the order of signaling events, the activated homotypic and heterotypic adhesion strengths, and the cadherin induction constants. In these rulesets (described in Figures 6–10), the GFP/Ncad phenotype and mCherry/Pcad phenotype can take on different homotypic adhesion strengths and induction constants (e.g., [*c*_*a*_, *P*_*a*_] were assigned to GFP/Ncad cells, and [*c*_*b*_, *P*_*b*_] were assigned to mCherry/Pcad cells). Each ruleset and its associated observations are explained in detail in *Results*.

### k-means clustering and tSNE dimensionality reduction

k-means clustering of the parameter space exploration results was conducted using the scikit-learn library in Python (71). The optimal number of clusters for each clustering problem was determined by plotting the within-cluster standard deviation for k-means clustering using two to fifteen clusters. This analysis was used to determine the minimal number of clusters that reduced within-group standard deviation. tSNE dimensionality reduction was conducted using the same Python library.

### Statistics

Statistical comparisons for continuous data were made by a one-way Kruskal-Wallis test followed by Dunn’s post-hoc analysis. This test was specifically applied when comparing fractional areas for *in vitro* spheroids and in predicted ABM spheroids (Figure 2). Statistical comparisons for qualitative metrics (e.g., core cell type) and discrete metrics (e.g., number of clusters) were made using a Chi-Squared test with *p* < 0.05 considered significant for all tests.

### Code Availability

Code for the 2D and 3D ABM, evolutionary algorithm, quantification of model and *in vitro* outputs, and k-means clustering of parameter space explorations are available here: https://github.com/nikita-sivakumar/multicell.

## ACKNOWLEDGEMENTS

The authors gratefully acknowledge funding from the University of Virginia Engineering in Medicine Program, the University of Virginia Center for Advanced Biomanufacturing, the Arnold and Mabel Beckman Foundation, and the National Science Foundation (1700687 to MJL). The authors are grateful to Dr. Wendell Lim (University of California San Diego) for providing engineered L929 cell lines and to Dr. Satoshi Toda and Dr. Nisha G. Sosale for technical discussions.

